# Identify Regulatory eQTLs by Multiome Sequencing in Prostate Single Cells

**DOI:** 10.1101/2024.06.19.599704

**Authors:** Yijun Tian, Lang Wu, Chang-Ching Huang, Liang Wang

## Abstract

While genome-wide association studies and expression quantitative trait loci (eQTL) analysis have made significant progress in identifying noncoding variants associated with prostate cancer risk and bulk tissue transcriptome changes, the regulatory effect of these genetic elements on gene expression remains largely unknown. Recent developments in single-cell sequencing have made it possible to perform ATAC-seq and RNA-seq profiling simultaneously to capture functional associations between chromatin accessibility and gene expression. In this study, we tested our hypothesis that this multiome single-cell approach allows for mapping regulatory elements and their target genes at prostate cancer risk loci. We applied a 10X Multiome ATAC + Gene Expression platform to encapsulate Tn5 transposase-tagged nuclei from multiple prostate cell lines for a total of 65,501 high quality single cells from RWPE1, RWPE2, PrEC, BPH1, DU145, PC3, 22Rv1 and LNCaP cell lines. To address data sparsity commonly seen in the single-cell sequencing, we performed targeted sequencing to enrich sequencing data at prostate cancer risk loci involving 2,730 candidate germline variants and 273 associated genes. Although not increasing the number of captured cells, the targeted multiome data did improve eQTL gene expression abundance by about 20% and chromatin accessibility abundance by about 5%. Based on this multiomic profiling, we further associated RNA expression alterations with chromatin accessibility of germline variants at single cell levels. Cross validation analysis showed high overlaps between the multiome associations and the bulk eQTL findings from GTEx prostate cohort. We found that about 20% of GTEx eQTLs were covered within the significant multiome associations (*p*-value ≤ 0.05, gene abundance percentage ≥ 5%), and roughly 10% of the multiome associations could be identified by significant GTEx eQTLs. We also analyzed accessible regions with available heterozygous SNP reads and observed more frequent association in genomic regions with allelically accessible variants (*p* = 0.0055). Among these findings were previously reported regulatory variants including rs60464856-*RUVBL1 (*multiome *p*-value = 0.0099 in BPH1) and rs7247241-*SPINT2 (*multiome *p*-value = 0.0002- 0.0004 in 22Rv1). We also functionally validated a new regulatory SNP and its target gene rs2474694-*VPS53 (*multiome *p*-value = 0.00956 in BPH1 and 0.00625 in DU145) by reporter assay and SILAC proteomics sequencing. Taken together, our data demonstrated the feasibility of the multiome single-cell approach for identifying regulatory SNPs and their regulated genes.

## Introduction

Although genome-wide association study (GWAS) has been highly productive in finding disease risk loci, only a small portion of these single nucleotide polymorphisms (SNPs) have been functionally characterized (Gallagher and Chen-Plotkin 2018; Farashi et al. 2019; van Bree et al. 2022). Thus far, as most risk-SNPs are found in noncoding regions of the genome, it is believed that many of them (or their closely linked SNPs) would alter regulatory element activities and quantitatively change gene expression rather than directly mutate protein sequences (Ernst et al. 2011; Maurano et al. 2012; Trynka et al. 2013; Weighill et al. 2022). To dissect functional variants and their target genes, expression Quantitative Trait Loci (eQTL) analysis has been commonly used to evaluate cis-regulatory variants tuning gene expression (Thibodeau et al. 2015). <u>Despite contributing to a better understanding of the biological significance of risk SNPs,</u> <u>the eQTL analysis has its technical challenges, including</u> 1) bulk tissue-based eQTL data are from a mixture of multiple cell types, which may dilute the signals produced by cells of interest and thus reduce sensitivity (Geeleher et al. 2018), 2) eQTL analysis often calls significant associations of a single target gene with multiple SNPs showing highly linkage disequilibrium (LD), leaving it difficult to dissect causal SNPs, and 3) eQTL analysis requires a large sample size (often at least hundreds of individuals) to detect weak effects of SNPs on gene expression.

To address these limitations, single-cell RNA sequencing (scRNA-seq) has recently been used in eQTL analysis with multiplexed patient samples to detect genotype-to-gene expression associations (Auerbach et al. 2021; Eraslan et al. 2021; Neavin et al. 2021). With the genotype information captured in the RNA-seq and the cell type classified from the single-cell transcriptome, this method focuses on resolving the cell-type specific signal and provides a powerful annotation to the bulk tissue eQTL findings. However, since the technology did not quantify the regulatory potential of the target variants, it can not decide the causality of the SNPs. Moreover, the sparsity nature of scRNA-seq data poses a challenge to the statistical power of between-cell association tests, restricting the broader application of this method (Dong et al. 2021; Liu et al. 2023). Meanwhile, to uncover the functional aspect of genomic DNA, the Assay for Transposase Accessible Chromatin with high-throughput sequencing (ATAC-seq) has been extensively applied for mapping regulatory elements at genome-wide scale (Buenrostro et al. 2015; Vu et al. 2023). With development of feature barcoding technology, multiomics sequencing has successfully combined ATAC-seq with RNA-seq at the single-cell levels (Ma et al. 2020), which provides a unifying workframe for dissecting functional variations that could cause gene expression changes. Additionally, to address sparsity issue of single cell sequencing, target hybridization capture has been proposed to enrich the information at regions of interest, which may improve single-cell data quality (Pokhilko et al. 2021).

To functionally characterize eQTL findings, we performed multiome single-cell sequencing in a total of 65,501 cells, originating from multiple human prostate cell lines. We further performed target hybridization capture using the multiome single-cell sequencing libraries to gain deeper insights into the eQTL catalog identified from bulk sequencing cohorts. With the information gained, we sought to develop an analytical workflow to detect epigenetic impact of germline variations on cis-regulation of gene expression.

## Results

### Multiome single-cell ATAC-seq captured transcriptional factor footprint and allele specific accessibility

To simultaneously capture the chromatin accessibility and gene expression profiling, we performed multiome single-cell sequencing in eight prostate cell lines, including RWPE1, RWPE2, PrEC, BPH1, DU145, PC3, 22Rv1 and LNCaP. The details about the data coverage and the quality control information can be accessed from Methods and Materials section and **Supplementary Table 1**. Combined analysis identified a total of 65,501 single-cell events with high quality multiomic data for all cell lines. Since single-cell ATAC-seq library captures accessible chromatin fragments by transposase insert, as revealed by the fragment size distribution (**Fig. S1A**), a majority of the ATAC-seq read pairs captures nucleosome-free regions (0∼150bp). The ATAC-seq UMAP demonstrated a clear separation among the eight separate samples (**Fig. S1B**). Notably, the androgen-sensitive cell LNCaP showed two separate clusters between DHT and methanol treatment, while the androgen-independent cell 22Rv1 displays two overlayed clusters (**Fig. S1B**). To characterize the accessible chromatin regions related to transcription factor binding, we profiled Tn5 insertion signals at 633 motifs of JASPAR transcription factor collection. This analysis revealed significant enrichment of open chromatins at transcription factor binding sites. For instance, LNCaP and 22Rv1 cells exposed to DHT exhibited higher chromatin accessibilities at the androgen receptor binding sites than those treated with methanol only (**Fig. S1C**). Moreover, surrounding the TP53 binding motif, prostate cell lines with benign origins (PrEC, BPH1, RWPE1 and RWPE2) tended to show higher accessibilities than those originating from prostate malignancies, supporting a loss-of-functional tumor suppressor TP53 in prostate cancer (**Fig. S1D**). Furthermore, we observed a distinct footprint nearby FOXA1 (**Fig. S1E**) and NRF2 (**Fig. S1F**) motifs in the genome scale, suggesting various function variability driven by these transcription factors. To investigate whether the allele-specific chromatin accessibilities will be reflected as fragment size difference, we examined the correlations between allele depth ratio and allelic fragment size ratio (ratio calculated from median sizes of the fragments carrying either reference or alternative allele), and found that the alleles with higher read depth tended to be present in smaller fragments (**Fig. S1G, S1H**). Consistently, alleles with higher depth tended to be present in fragments with higher nucleosome-free proportions (**Fig. S1I, S1J**).

### Multiome single-cell RNA-seq characterized HALLMARK pathway alteration and aneuploidy-related transcriptome change

Since the multiome single-cell RNA-seq applies 3’ gene expression barcoding technology, the majority of coverage was enriched in the transcription ending sites of known human genes (**Fig. S2A**). Consistent with the ATAC-seq UMAP, the RNA-seq cluster also showed similar separation in 22Rv1 and LNCaP cells with or without DHT treatment (**Fig. S2B**). To systematically characterize transcriptome alterations in known pathways, we performed single-cell gene set enrichment analysis in Molecular Signature Database (MSigDB). Among the 50 HALLMARK gene sets, we found consistent androgen response pathway (Liberzon et al. 2015) activation in the DHT treated 22Rv1 and LNCaP cells (**Fig. S2C**). Moreover, we found an activation of KRAS signaling (Bello et al. 1997) in RWPE2 cells when compared with its parental line RWPE1, demonstrating the overexpression status of KRAS in RWPE2 in nature (**Fig. S2C**). We also performed similar enrichment analysis with positional gene sets, and observed variable sectorized transcriptome changes across different cell lines (**Fig. S2D**), which reflected gross genomic abnormalities.

### Target enrichment enhanced data quality for integrative analysis in multiome single-cell sequencing

One drawback of single-cell sequencing is data sparsity, which has limited the downstream statistical analysis. To increase sequencing depth at the genomic regions of interest, we performed a targeted enrichment in the regions covering risk SNPs and their associated genes using the single-cell libraries (**Fig. 1A**). For the RNA-seq panel, we designed hybridization probes targeting 273 genes based on multiple prostate eQTL databases, include MAYO prostate tissue, TCGA Prostate Adenocarcinoma and GTEx prostate cohorts (**Fig. 1B**). For the ATAC-seq panel, we included 2,730 eQTL SNPs located in the single-cell ATAC-seq peak regions (**Fig. 1C**). The data from the target sequencing showed roughly the same cell number as the corresponding parental sample (**Fig. 1D**). To compare the between-cell transcriptome differences related to the effect of allelic variants, we first quantitated the Tn5 insertion to distinguish the accessibility of certain SNPs in the cell. Specifically, for each SNP locus, if at least one Tn5 insertion was found in the cell, the cell was assigned to the “cut” group. Otherwise, if no Tn5 insertion is found, the cell is assigned to “no cut” group” (**Fig. 1E**). We then tallied the genes and SNPs informed in each sample and observed an increased number of cells with informative eQTL information in after-enrichment libraries (**Fig. 1F, 1G**). More quantitatively, we calculated the abundance of gene expression cell matrix and accessible chromatin cell matrix, which represented the proportion of non-missing value in each sample. While the abundance remained low, the additional target sequencing improved the eQTL gene expression abundance by 20% (**Fig. 1H**) and the eQTL accessibile chromatin abundance by 5% (**Fig. 1I**).

**Figure 1.**
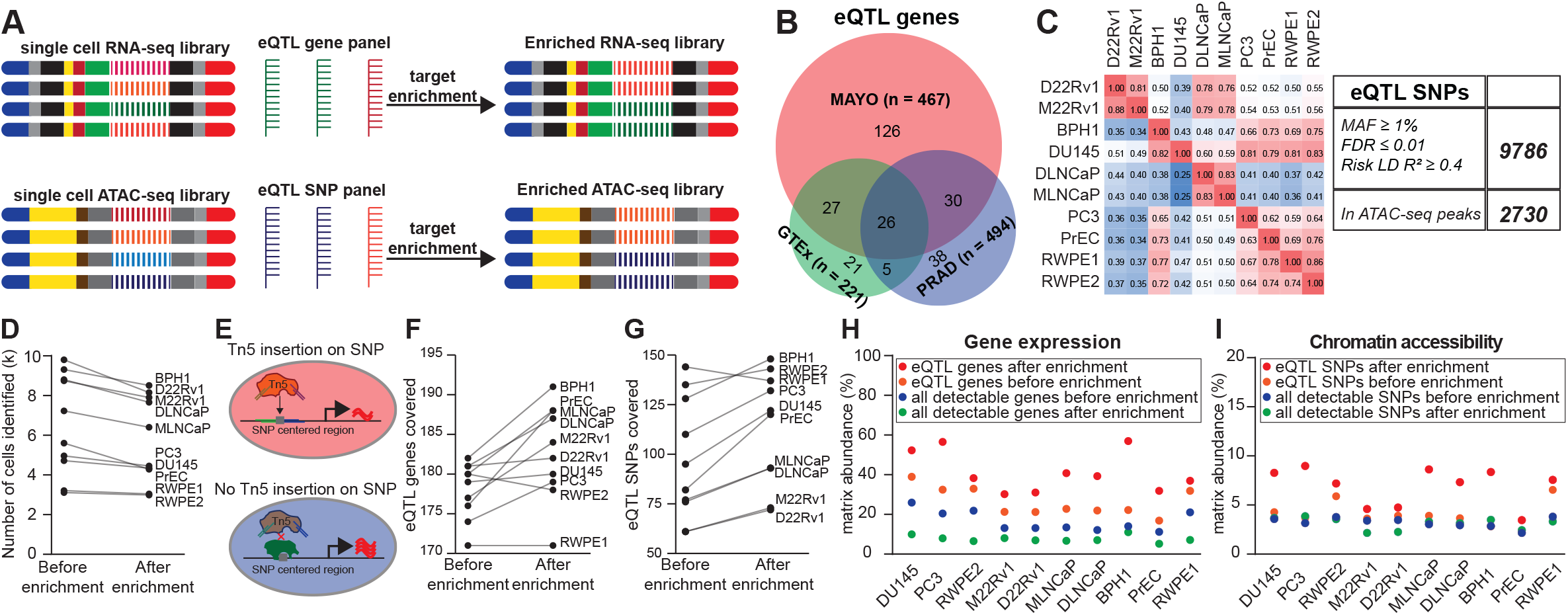
**A.** Experimental design combining multiome single-cell sequencing and targeted hybridization capture to enrich scATAC-seq and scRNA-seq signals for regulatory element analysis. **B.** Venn diagram of targeting genes previously identified in prostate cohorts, including GTEx prostate tissue, MAYO normal prostate and TCGA prostate adenocarcinoma. Numbers in circle indicate unique numbers of genes across each section, while numbers in bracket indicate cohort sample sizes. **C.** Pairwised intersecting heatmap for peak overlapping status in ATAC-seq data of all prostate cells, and the eQTL SNP selection criteria. Numbers in each heatmap cell indicate the intersected peak portion to the collection of the vertical sample. **D.** Comparison of the cell numbers identified covered in the single-cell multiome libraries before and after targeted capture. **E.** Schematic demonstration about the strategy to determine whether a cell carries a Tn5 insertion at certain SNP. **F.** Comparison of the eQTL genes covered in the single-cell multiome libraries before and after targeted capture. **G.** Comparison of the eQTL SNPs covered in the single-cell multiome libraries before and after targeted capture. **H.** Gene expression matrix abundance in each single-cell multiome sequencing library. **I.** ATAC-seq derived SNP signal matrix abundance in each single-cell multiome sequencing library.

### Multiome single-cell associations were partially overlapped with eQTL findings in prostate tissues

To identify target genes influenced by the accessible state of certain SNPs, we performed differential expression analysis between “cut” and “no cut” cell classes. We first characterized the overlap between prostate cell multiome associations and GTEx prostate tissue eQTL signals. This analysis showed that about 20% of GTEx eQTLs were covered within the significant multiome associations (p-value ≤ 0.05 and with tested genes detected in at least 5% cells) (**Fig. 2A**), supporting the notion that only a fraction of the eQTL SNPs are regulatory to the target genes. Additionally, 10% of the multiome significant associations could be identified by GTEx eQTLs (**Fig. 2B**), indicating that the prostate-specific regulatory effect could be interfered with by multiple cell type signals in bulk eQTL cohorts. Additionally, we evaluated the consistency between different cell lines. Among the 6,201 significant associations, we identified 1,247 multiome associations (roughly 20% of all findings) that were shared by at least 2 cell lines. We further characterized the overlapped findings within subgroups according to the cell line origins and conditions, such as normal prostate, prostate cancer and androgen treatment status (**Fig S3A-S3C**). Specifically, we identified 111, 79, and 71 risk-related associations from the shared findings in normal prostate (37.2%), prostate cancer (43.4%) and androgen treatment (51.4%) subgroup, respectively. Although the current multiome association analysis sorely examined the regulatory potential of the chromatin accessibility regardless of the allelic impact, we wondered if the association signals attributed to the allele difference. To this end, we compared the proportions of significant associations between homozygote and heterozygote SNPs and found no significant difference in regard to the genotypes (**Fig. 2C**). However, when comparing the significant associations between equally and allelically accessible loci, we found a higher proportion of significant associations in those variants showing allelic accessibility (p=0.0055, **Fig. 2D**).

**Figure 2.**
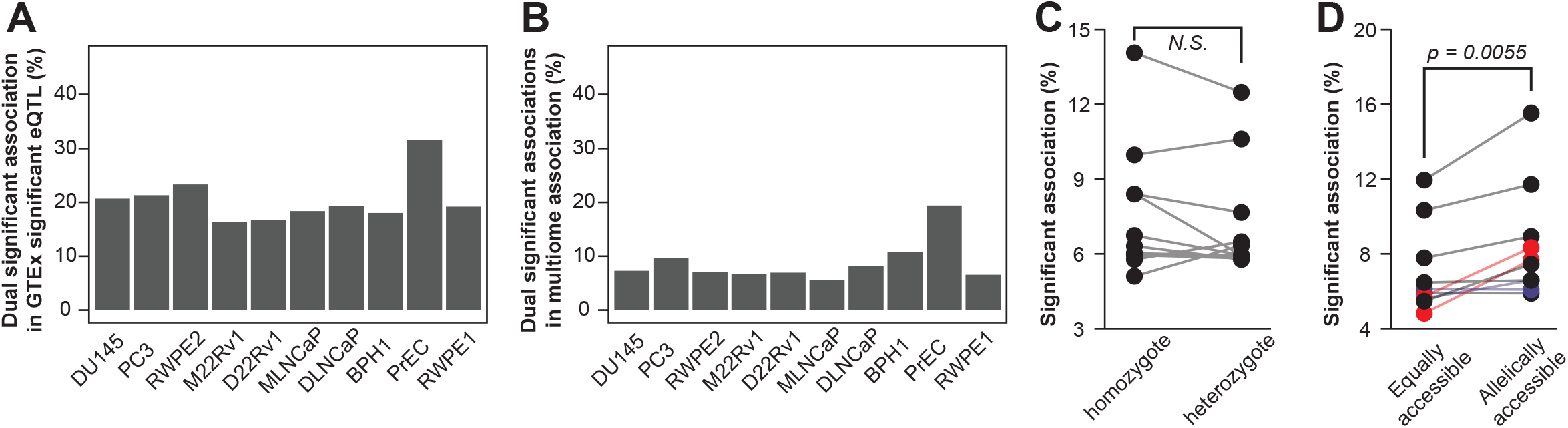
**A-B.** Proportion of dual significant (significant in both single-cell multiome sequencing and GTEx prostate cohort) SNP-to-Gene association to GTEx significant eQTLs (**A**), or to significant single-cell multiome associations (**B**). **C.** Comparison of the significant multiome association proportions with homozygote or heterozygote genotypes in all single-cell sequencing samples. **D.** Comparison of the significant multiome association proportions with equally or allelically accessible variant in each cell line sample.

### Multiome single-cell association test identified rs2474694-VPS53 functional locus in prostate cells

Among the multiome associations, we further evaluated a prostate cancer GWAS risk locus in BPH1, DU145 and PC3 cell lines. As reported by the FinnGen study (Kurki et al. 2023), the rs2474694 A allele contributed to higher risk of prostate cancer than G allele (p = 3.22e-9, OR = 1.09, 95% CI: 1.08,1.11). In BPH1 and DU145 multiome data, we found that the cells with “cut” at rs2474694 locus showed significantly higher VPS53 expression than cells with “no cut” (**Fig. 3A**). Using luciferase reporter assay, we identified that rs2474694 G exhibited consistently stronger transcription activity than A allele in BPH1, DU145 and PC3 cells (**Fig. 3B**). Interestingly, among the VPS53 eQTL SNPs, we observed the best p-value for rs2474694 in GTEx cohort (**Fig. 3C**). To understand which protein factor binds on the rs2474694 allelically, we performed Mass Spectrometry proteomics with SILAC labeled nuclear protein, and found various protein kinase, GTPase and zinc finger proteins binding differentially to the rs2474694 alleles, such as CDC42, RHOA/RHOB and ZNF668 (**Fig. 3D**). In ANANASTRA (Annotation and enrichment Analysis of Allele-Specific Transcription factor binding at SNPs) database, we also identified multiple transcription factor that exhibited allele-specific signals in published ChIP-seq experiments (Abramov et al. 2021) (**Fig. 3E**). Lastly, from a CRISPRi screening database (Tian et al. 2023), the rs2474694 locus showed an essential role in BPH1 and PC3 cell proliferation, which demonstrated its pivotal role in driving prostate cancer risk (**Fig. 3F**).

**Figure 3.**
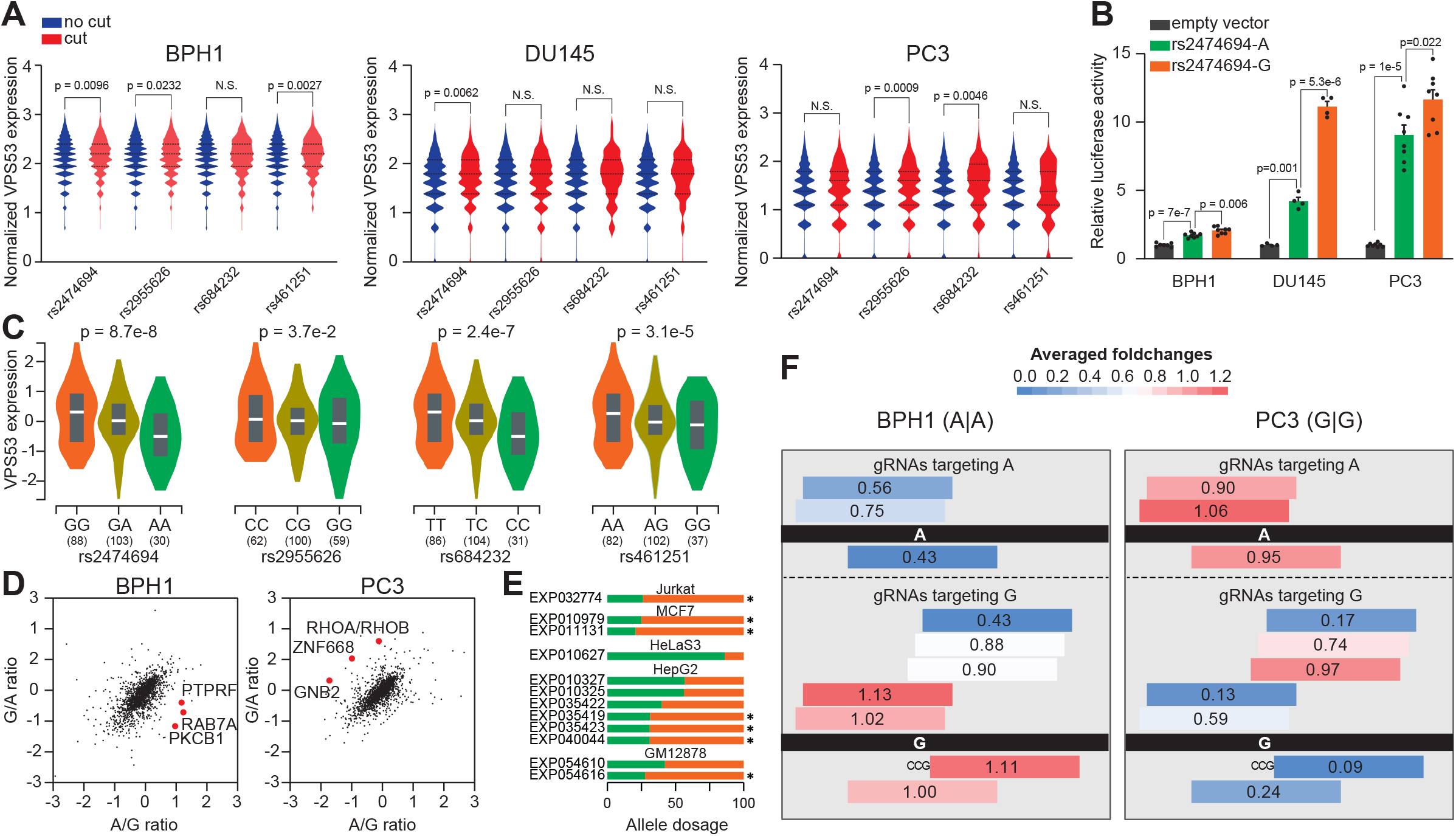
**A.** Violin plot of single-cell VPS53 gene expression between cells with or without cut at specific SNP locus. **B.** Luciferase reporter assay comparing rs2474694 allele transcription activities in BPH1 and PC3 cells. Individual dot indicates an independent biological replicate. **C.** Violin plot of VPS53 gene expression in bulk prostate samples with various genotypes. **D.** Allele specific protein binding profiling at rs2474694 locus by SILAC-based proteomics analysis. **E.** Allele specific TF binding evidence of other human cell lines identified in ANANASTRA database. **F.** CRISPRi screening proliferation fold change of rs2474694 gRNA in BPH1 and PC3 cells. The black bar indicates rs2474694 locus in the germline, and colored strips above or below the bar indicate gRNA binding Crick or Watson strands of the germline DNA. Numbers in the center show the average fold change for two screening replicates.

Additionally, the multiomic association analysis also identified another functional locus rs60464856-*RUVBL1*. This SNP-gene pair showed the best statistical significance across multiple LD SNPs (**Fig. S4A**) and positive regulatory potential in a previous publication (Tian et al. 2023). For another previously validated functional SNP rs7247241 (Tian et al. 2022), we observed its multiome association significance for the SPINT2 gene expression changes in 22Rv1 cells. Interestingly, another linked eQTL SNP rs8100395 showed much weaker multiome significance in 22Rv1 cells. (**Fig. S4B**).

## Discussion

To date, the GWAS and eQTL findings present an equal number of opportunities and challenges to advance our understanding of prostate cancer etiology. Theoretically, each PCa risk locus and its related eQTLs are associated with underlying mechanisms contributing to PCa incidence. However, establishing causality between the variant and gene expression is highly challenging due to weak regulatory effects resulting from single DNA base alterations, which necessitate a clear-signal environment for observation. The prostate cell line provides such an environment because the cell lines originate from a monoclonal progenitor and exhibit similar epigenetic modifications and transcriptome profiling. These advantages make cell lines well-suited for assay and pipeline development to detect functional variants that are capable of changing the chromatin accessibility and gene expression. Therefore, applying the multiome sequencing in cell lines is feasible for functional variant analysis. However, there remains a critical drawback of single-cell technology, which is the data sparsity. Targeted single-cell RNA-seq has been proven to greatly improve biological information (Pokhilko et al. 2021). Similar to the findings, the targeted enrichment in this study also increased the expression abundance of the eQTL gene from the multiome single-cell RNA-seq library. In contrast, the improvement of accessible chromatin abundance in the single-cell ATAC-seq data is relatively small. We speculate that this difference largely arises from starting template abundance variations between DNA and RNA. It is estimated that the amount of RNA per cell is around 10-30 pg total RNA, and each diploid human cell contains approximately 6pg genomic DNA (Abramov et al. 2021). With a more abundant template amount, gene expression will benefit more from the targeted enrichment process. For scATAC-seq libraries, although the improvements were limited, the enriched data tended to receive more coverage at selected eQTL SNPs, which was also a significant advantage for direct comparison to known eQTL findings.

Our study showed a strong signal overlap between multiome tests and eQTL analysis. Particularly, the multiome association from the PrEC cell line showed the highest fraction of overlap signals with bulk prostate tissue eQTLs. Since PrEC cell line is derived from benign prostate tissue, it is not surprising to see the higher signal overlap between this cell line and bulk prostate tissues. Nevertheless, this study provided a comprehensive association list between accessible chromatins and cis-gene expression. This list will significantly enrich the functional annotation of current eQTL databases, which are largely limited in the single-cell eQTL studies (van der Wijst et al. 2018). Additionally, the current multiome pipeline did not directly examine the functional effect of SNP alleles on the gene expression. With an increased coverage at selected eQTL loci, we reasoned that the multiome analytical pipeline could be expanded to evaluate the allele accessibility and downstream gene expression. Clearly, the data from this study support this application.

Among our findings, we highlighted our finding that showed higher *VPS53* expression in the cells with a Tn5 cut near the rs2474694, which provided functional support for the eQTL association found in the bulk sequencing cohort. Notably, a Tn5 cleavage near a SNP indicates an accessible state of the genomic DNA capable of interacting with a transcription factor, indicating a stronger regulatory impact contributing to a different gene expression in that cell. The reporter assay, SILAC proteomics and the published ChIP-seq data consistently demonstrated the allelic difference, further supporting the validity of our multiome pipeline in identifying functional SNPs. Interestingly, from the multiome association findings, we confirmed another 2 regulatory variants, which had been functionally characterized in previous studies. The rs60464856, identified from BPH1 multiome data, had been proven to be capable of regulating *RUVBL1* gene expression in *BPH1* base editing experiment (Tian et al. 2023). The rs7247241, identified from 22Rv1 multiome data, had been found to be able to change proximal genome methylation in 22Rv1 cells (Tian et al. 2022). More interestingly, compared to the adjacent LD SNP rs8100395, the multiome method demonstrated stronger association for rs7247241, further supporting its potential of identifying regulatory variants for functional eQTL characterization. Based on these observations, we anticipate broader usage of the multiome single-cell sequencing technology to investigate the functional implications of noncoding variants.

Although the multiome approach is promising, the current study has its own pitfalls. First, we applied targeted hybridization capture to enrich read counts at the regions of interest. Although the additional enrichment substantially improved the gene expression in the multiome associations, there was a limited improvement in chromatin accessibility. The scATAC-seq data abundance could be enhanced with a better library preparation protocol, including but not limited to development of high efficiency library preparation protocol, application of pre-amplified library for targeted hybridization, and capture of more single cells. Second, due to the technical constraints, we were only able to obtain high quality multiome data using prostate-derived cell line. To gain native insight of prostate-specific regulatory network, it is necessary to expand the current methodology to prostate tissues and perform multiome association analysis within each cell type. Furthermore, owing to lack of multiplex approach, the current processing cost of the multiome method is expensive, which poses a significant challenge to test this approach in large cohort. To obtain a comprehensive atlas of those functional variants, cost-effective approach to pool nuclei from different individuals is warranted to increase genetic diversity.

Taken together, by combining existing 10X multiome protocol with target hybridizatioin method, we developed an innovative workframe to dissect regulatory SNPs that can change gene expression profiling, and provided functional insight into eQTL association at prostate cancer risk loci. This cell lines-based study supports application of this workframe in the functional characterization of disease risk..

## Methods

### Cell culture and reagents

DU145 (CVCL_0105), PC3 (RRID: CVCL_0035), 22Rv1 (RRID: CVCL_1045), LNCaP (RRID: CVCL_0395), RWPE-1 (RRID: CVCL_3791), RWPE-2 (RRID: CVCL_3792) and PrEC (RRID: CVCL_0061) cells were obtained from the ATCC. BPH1 (RRID: CVCL_1091) cells were purchased from Sigma-Aldrich. All cell lines were examined for mycoplasma contamination with Venor GeM Mycoplasma Detection Kit (Sigma-Aldrich). Unless specified otherwise, all cell culture reagents were obtained from ThermoFisher Scientific. BPH1, DU145 and PC3 cells were grown in RPMI1640 medium supplemented with 10% fetal bovine serum (FBS). RWPE1 and RWPE2 cells were grown in Keratinocyte Serum-Free Medium. For androgen starvation, 22Rv1 and LNCaP cells were grown in phenol red free RPMI 1640 media supplemented with 10% charcoal stripped fetal bovine serum (FBS) for 16 hours. After starvation, 5 nM Dihydrotestosterone or same volume of solvent were added to the the media to treat the cells for total of 16 hours. The cells were dissociated into single-cell suspension with TrypLE Expression Enzyme, washed with ice-cold PBS twice and strained with a 40 micron flowmi strainer before the single-cell encaptualization.

### Multiome single-cell sequencing

The multiome libraries were prepared according to the manufacturer’s protocol. Briefly, nuclei were isolated from the dissociated cells and the open chromatin regions were tagged with Tn5 transposase. The tagmentated nuclei of each cell line was then applied to one lane of the Chromium Next GEM Chip J for GEM generation and molecule barcoding. After GEM incubation and library construction, each ATAC-seq read was labeled with 10x cell identity barcodes, while each RNA-seq read was captured by paired barcodes with an Unique Molecular Identifier (UMI). For raw multiome library capture, we aimed 8,000-10,000 nuclei for each sample and 25,000 scATAC-seq, and 20,000 scRNA-seq reads per cell to ensure the data coverage. For ATAC-seq, we kept cells with good transcription starting site (TSS) enrichment (>1) and nucleosome signals (<2), as well as enough raw per-cell ATAC, read count (>1000). For gene expression data, we kept cells with low mitochondria gene percentage (<50%) and enough per-cell RNA read count (>200). We aimed for a similar total read count for each targeted enriched library for better comparisons. The summary of the Cellranger-ARC count can be found in **Supplementary Table 1**.

### Targeted sequencing by hybridization capture

To gain insights about the functional eQTL findings, we sought to use hybridization capture to improve the data quality. We designed two target enrichment panels to capture the gene expression and the chromatin accessbility signals in related to previous eQTL findings.

For the gene expression panel, we selected 273 eQTL genes (c2) from 3 prostate eQTL datasets, including MAYO, GTEx and TCGA adenocarcinoma cohorts. We also included 14 internal control gene (c1) in the RNA panel to benefit the normalization process. For eQTL genes, we designed as many probes as possible based on the cDNA and the 3’UTR sequences of each isoform. Since the internal control genes are generally with high RNA abundance, we choose to design 1 to 2 high quality probes to cover only the exon with the highest expression from the raw sequencing data. As a result, we designed 16,669 RNA probes for the gene expression panels to cover 2,000,280 bp.

For the ATAC-seq panel, we selected 9,786 SNPs within the MAYO prostate cohort based on the below criteria: 1) with Minor allele frequency (MAF) in the Caucasian population ≥ 1%; 2) with False discovery rate (FDR) of any eQTL association ≤ 0.1; 3) with R^2^ of linkage equilibrium to any prostate cancer risk SNP ≥ 0.4. To increase the hybridization efficiency, we further focused on the open chromatin regions for probe design. Considering that the similarity of the chromatin accessibility regions between different prostate cell lines, we used the merged ATAC-seq peak region to further filter the SNPs to increase targeted capturing efficiency. As a result, we designed 2,120 probes to cover 2,730 SNPs, covering a total of 246,992 bp.

We used Standard Hyb and Wash Kit (v2) from Twist Bioscience to perform the targeted hybridization workflow, and chose NEXTFLEX® Universal Blockers to reduce non-specific hybridization from TruSeq (scRNA-seq) and Nextera (scATAC-seq) adapters. To perform multiplex hybridization capture, we measured concentration of the raw libraries with NEBNext® Library Quant Kit, and pooled equal molars of each library as a total of 2.5 ug input DNA. After the capture, we amplified 13 and 11 cycles for the scATAC-seq and the scRNA-seq library, respectively.

### Single-cell multiome sequencing data processing

To demultiplex the multiome samples and generate the FASTQ files for the single-cell quantification, we used Cellranger-ARC mkfastq to perform raw base calling from both ATAC and RNA flow cells. With the FASTQ files ready, we then used Cellranger-ARC count to perform sequencing alignment, cell barcode quantification, ATAC-seq peak calling, and RNA-seq gene expression quantification. After the counting process, we obtained the feature-barcoding matrices, the ATAC-seq fragment and peak information, and other intermediate files for downstream multiome association analysis. The summary of the Cellranger-ARC count can be found in **Supplementary Table 1**.

### Multiome association analysis

To discern the chromatin accessible states of each SNP using the ATAC-seq data, we devised a script (cutmat.sh) to extract the Tn5 insertion sites and generate a matrix that associates cells with SNPs, indicating whether a certain SNP region receives a Tn5 cleavage within the cell. More specifically, we retrieved the Tn5 insertion sites from ATAC-seq fragments by extending the coordinates of their ends by 5 bp. To intersect these Tn5 insertion sites with the SNP loci, we also extended each SNP coordinates by 20bp, considering that the length of most transcriptional factor motifs is 18-20 bp.

For gene expression data, we applied SCTransform (Hafemeister and Satija 2019) from Seurat (Hao et al. 2024) package to perform the normalization of count data from scRNA-seq information. To conduct the multiome association analysis, we compared the expressions of the genes located within 1 Mb window between the cells with and without a cut on each interest SNP. We utilized the FindMarkers function from Seurat (Hao et al. 2024) package to perform the comparison with multiple statistical methods, including the Student’s t-test (t), the ROC analysis, the Likelihood-ratio test (McDavid et al. 2013), the Wilcoxon Rank Sum test, the negative binomial generalized linear model, the Poisson generalized linear model, and logistic regression model. The statistical results for these methods can be found in **Supplementary Table 2**.

### Comparative benchmarking multiome association in prostate cell line with prostate GTEx eQTLs

To evaluate the overlapped proportion between the prostate cell line multiome association and the prostate GTEx findings, we defined significant findings for multiome associations as those with Student’s t-test p-value ≤ 0.05 and with tested genes detected in at least 5% cells for both “cut” and nocut cells, and defined significant findings for prostate GTEx eQTLs as those with nominal p-value ≤ 0.05. Measurements of the overlapped results were demonstrated in the below 2 × 2 table:

| | GTEx eQTL p-value $\leq 0.05$ | GTEx eQTL p-value $> 0.05$ |
| --- | --- | --- |
| Multiome p-value $\leq 0.05$ | **A** | **B** |
| Multiome p-value $> 0.05$ | **C** | **D** |

By this table, we calculated dual significant proportion in GTEx significant eQTL as 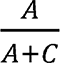 for each sample. Similarly, we calculated dual significant proportion in Multiome significant association as 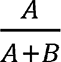 for each sample.

### Multiome association comparison by SNP genotype and allelic status

To identify the SNP genotype in the multiome findings, we extracted the genomic DNA and prepared whole genome sequencing library from each cell line using NEBNext® Ultra™ II DNA PCR-free Library Prep Kit. We then performed targeted capture using the DNA hybridization panel and used the WGS data to call variant genotypes. To estimate the allelically accessible dosage for the heterozygote variant, we developed a PERL script (ectInsBAM.pl) to extract the Tn5 insertion sequence from each unique scATAC-seq BAM alignment, and only used the insertion BAM for variant calling. After variant calling, we calculated the a odds ratio along with the 95% confidence interval for each heterozygote variant based on the reference and alternative allele dosage in WGS and ATAC-seq data. As a result, we distinguished the allele-specific chromatin accessibility for each SNP according to whether the confidence interval included odds ratio 1.

### Luciferase reporter assay

The cells were seeded into a 24-well plate. After 12 to 16 hours, 500 ng of pGL3 reporter plasmids were transfected to each well using Lipofectamine 3000. The media were replaced after transfection for 24 hours. After 48 hours of transfection, the cells were lysed for the luciferase assay according to Dual-Luciferase® Reporter Assay (E1960, Promega) protocol. The luminescence signals were measured with the GlowMax plate reader. After normalizing to Renilla luciferase readout, relative firefly luciferase activities driven by corresponding promoters were represented by luminescence unit fold changes.

### Allele-specific proteomics screening with stable isotope labeling by amino acid in cell culture (SILAC)

The BPH1 and PC3 cells were grown in SILAC RPMI 1640 medium (ThermoFisher 88365) for five passages before harvesting for nuclear protein extraction (Active Motif 40010). After confirming that heavy amino acid labeling efficiency reached 99.9%, the nuclear extracts were applied to the desalting spin column (ThermoFisher 89882) to remove excessive ions. The DNA baits harboring rs2474694 A and G alleles were produced according to a previous publication.^32^ For each binding reaction, 2 μg of purified DNA baits were conjugated to 25 μL Streptavidin Dynabeads (ThermoFisher 65001). The clean conjugated beads were incubated with 12.5 μL precleared nuclear protein at 4 degrees overnight. The incubated beads were washed five times and combined for two parallel quantitative mass spectrometry runs to get the allelic protein binding ratio.

### Source code and sequencing data accession

The source code generated in this study can be accessed through the github repository (https://github.com/Yijun-Tian/multiome). The raw data FASTQ file and the processed data matrix will become accessable through GEO accession GSE264518 upon formal publication.

## Acknowledgements

This study has been supported by National Institute of Health [R01CA250018 and R01CA212097, to L.Wang. and R01CA263494 to L.Wu]. This work has also been supported in part by the Molecular Genomics Core at the H. Lee Moffitt Cancer Center & Research Institute, a comprehensive cancer center designated by the National Cancer Institute and funded in part by Moffitt’s Cancer Center Support Grant (P30-CA076292). The funders had no role in study design, data collection and analysis, decision to publish, or manuscript preparation.

## Ethics approval and consent to participate

Not applicable

## Consent for publication

Not applicable

## Availability of data and materials

All data generated or analyzed in this study are included in this article or the supplemental information files.

## Authors’ Contributions

Conception and design: Y. Tian, L. Wang

Development of methodology: Y. Tian

Acquisition of data: Y. Tian

Analysis and interpretation of data: Y. Tian, L. Wu, C-C. Huang

Writing, review, and revision of the manuscript: Y. Tian, L. Wu, C-C. Huang, L. Wang

Study supervision: L. Wang

---

**Fig S1.**
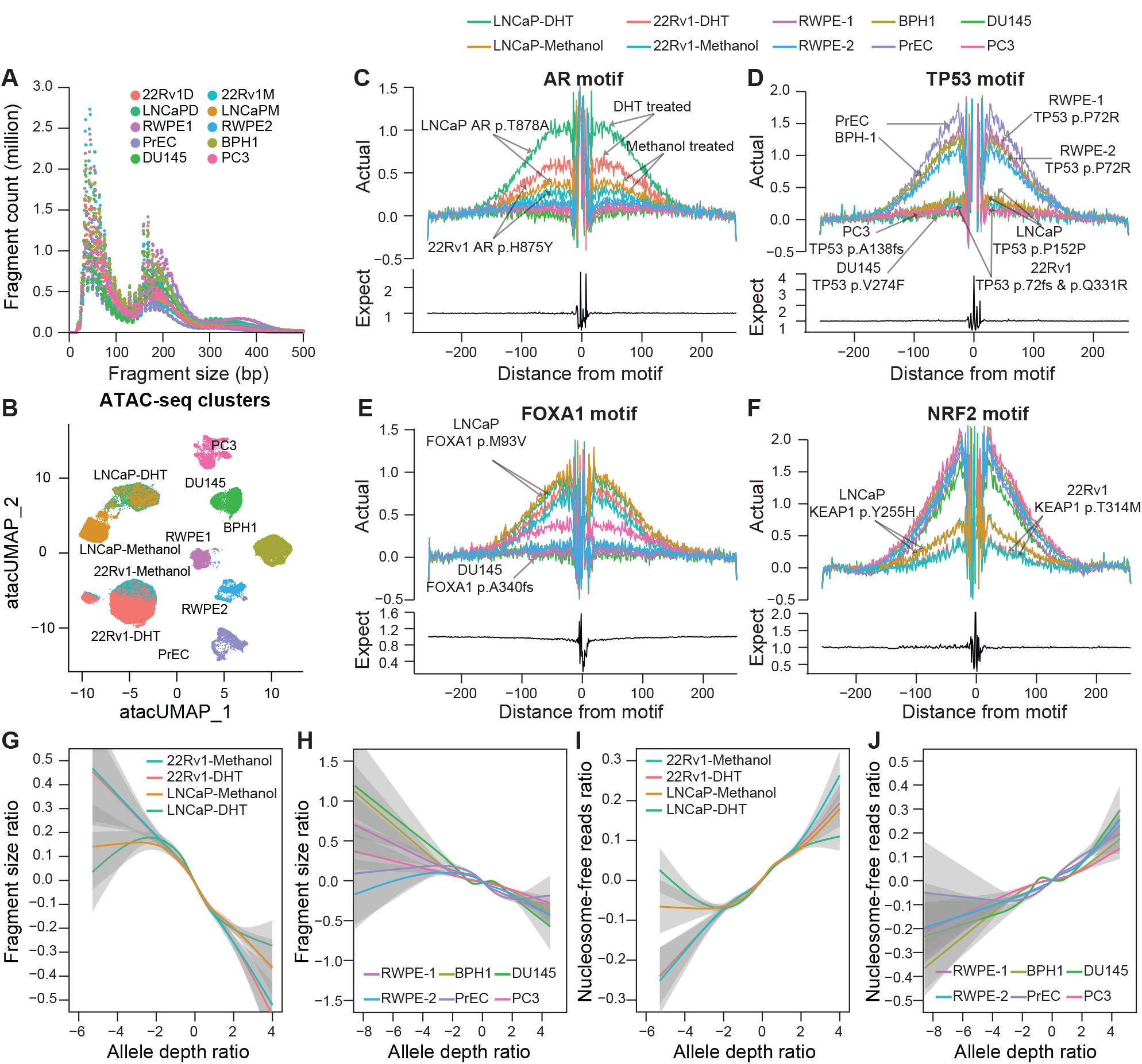

**Fig S2.**
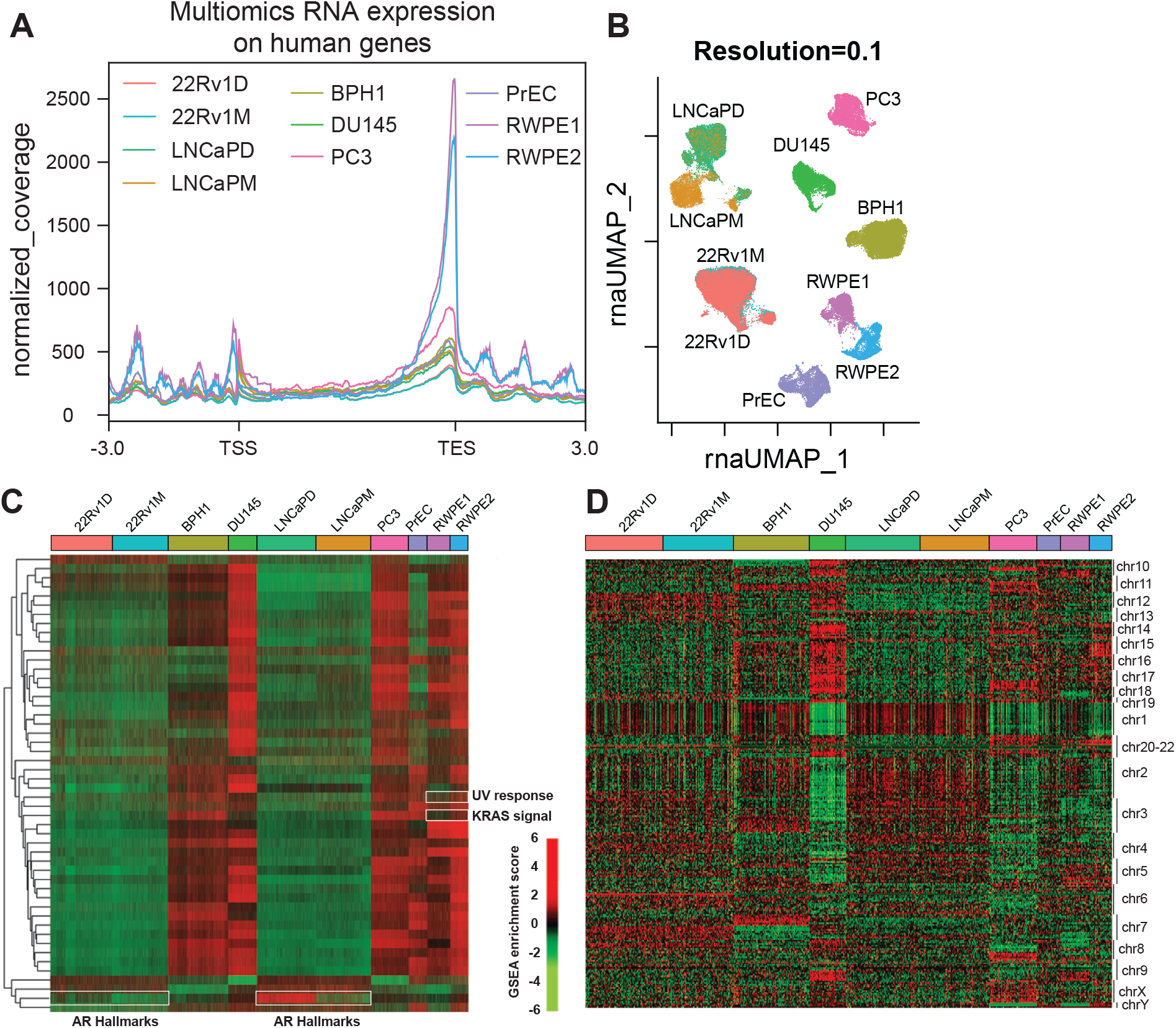

**Fig S3.**
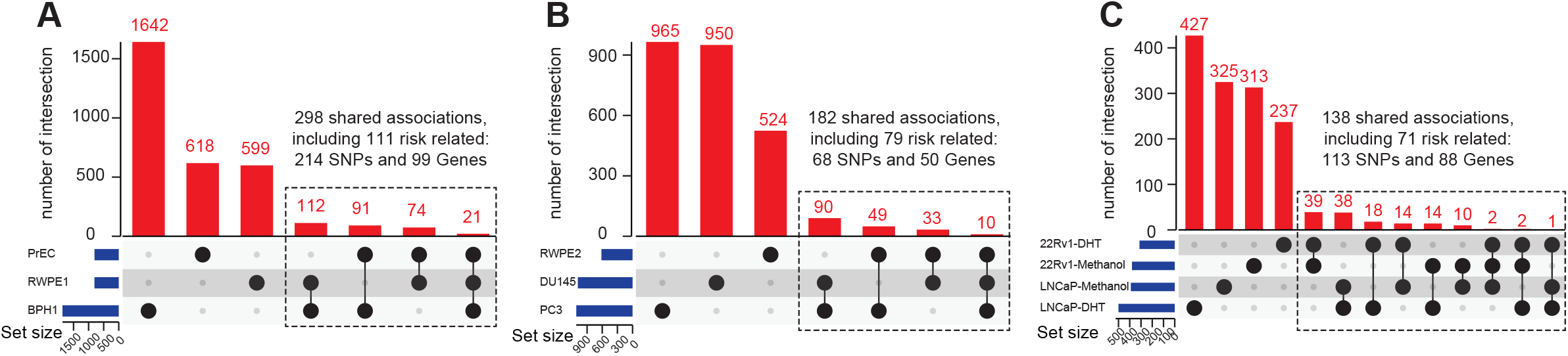

**Fig S4.**
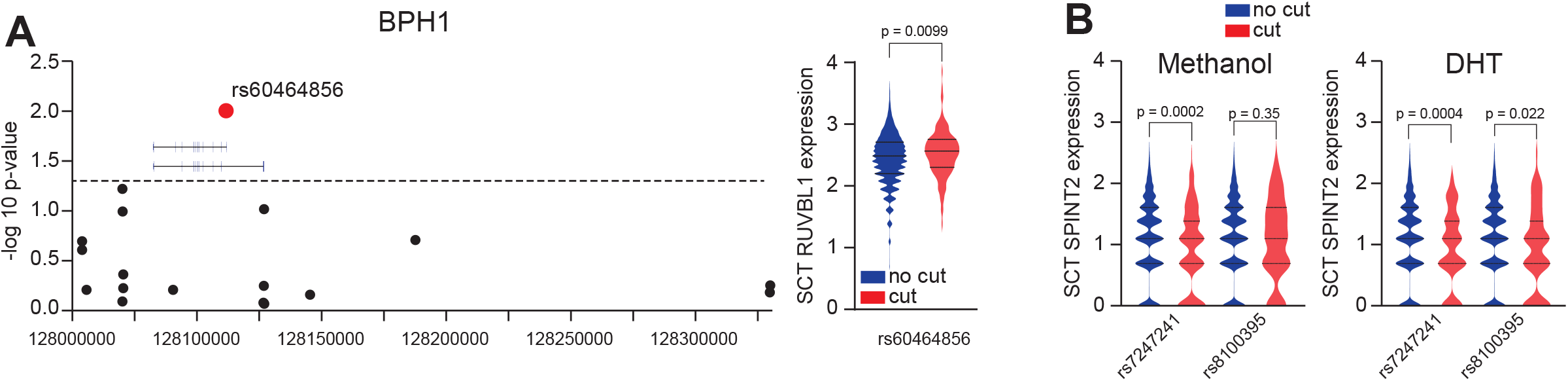

